# Discovery of novel enzybiotic candidates targeting human bacterial pathogens through large-scale viral-host profiling

**DOI:** 10.64898/2026.09.01.748673

**Authors:** Mateus B. Fiamenghi, Nikos C. Kyrpides

## Abstract

The rise of antibiotic-resistant bacteria demands alternative therapeutic strategies, with bacteriophage (phage) therapy and phage-derived enzybiotics emerging as promising approaches. However, identifying candidate phages against specific pathogens has historically been a bottleneck due to the need for cultivation methods to assess host range and lytic activity. Advances in metagenomic sequencing and the emergence of large-scale viral genome databases now provide an opportunity to accelerate this process computationally. Here, we present a large-scale mining of the MetaVR database to identify phages targeting human bacterial pathogens. By integrating direct host associations with CRISPR-spacer evidence, we linked 196,472 high-quality and complete viral genomes, representing 42,360 vOTUs, to 618 species of pathogenic and opportunistic bacteria. Functional enrichment analysis revealed distinct genomic signatures with viral lifestyle and host-range breadth: virulent phages were enriched in replication and structural functions, whereas temperate and broad host-range phages were enriched in anti-defense and regulatory modules. To characterize their lytic potential we annotated lysis-related protein families and their structural diversity, identifying 76 structurally novel lysis-associated proteins, including candidates targeting WHO priority pathogens. Focused analysis of endolysins revealed 592 structural clusters, with extensive sharing of endolysin repertoires among ESKAPE pathogens, suggesting candidates for broad-spectrum enzybiotic development. Selection analysis identified 167 endolysin families with sites under positive selection within functional domains, highlighting evolutionary diversification potentially associated with phage-host interactions. Together, our results establish a large-scale framework for connecting human bacterial pathogens to phages and their lytic machinery, providing a resource for prioritizing phage therapy and enzybiotic development.

**Importance:** Antibiotic resistance is a growing global health crisis, and phage therapy represents a promising alternative to conventional antibiotics. By leveraging a large-scale viral genomic database to computationally link almost 200,000 phage genomes to 618 human pathogen species, we substantially expand the catalog of known phage-pathogen interactions. Additionally, we characterize the lysis machinery encoded by these phages, including holins, spanins and endolysins uncovering structurally novel enzymes, associated with phages targeting WHO priority pathogens that could serve as starting points for enzybiotic development. We also reveal extensive sharing of endolysin repertoires among pathogen species, indicating opportunities for developing broad-spectrum enzybiotic strategies. By providing a systematic, genomics-driven framework for identifying and prioritizing phage-derived therapeutic candidates, this work translates large-scale viral genomic data into a resource for developing new therapeutic strategies against drug-resistant bacterial infections.

**Graphical Abstract:** 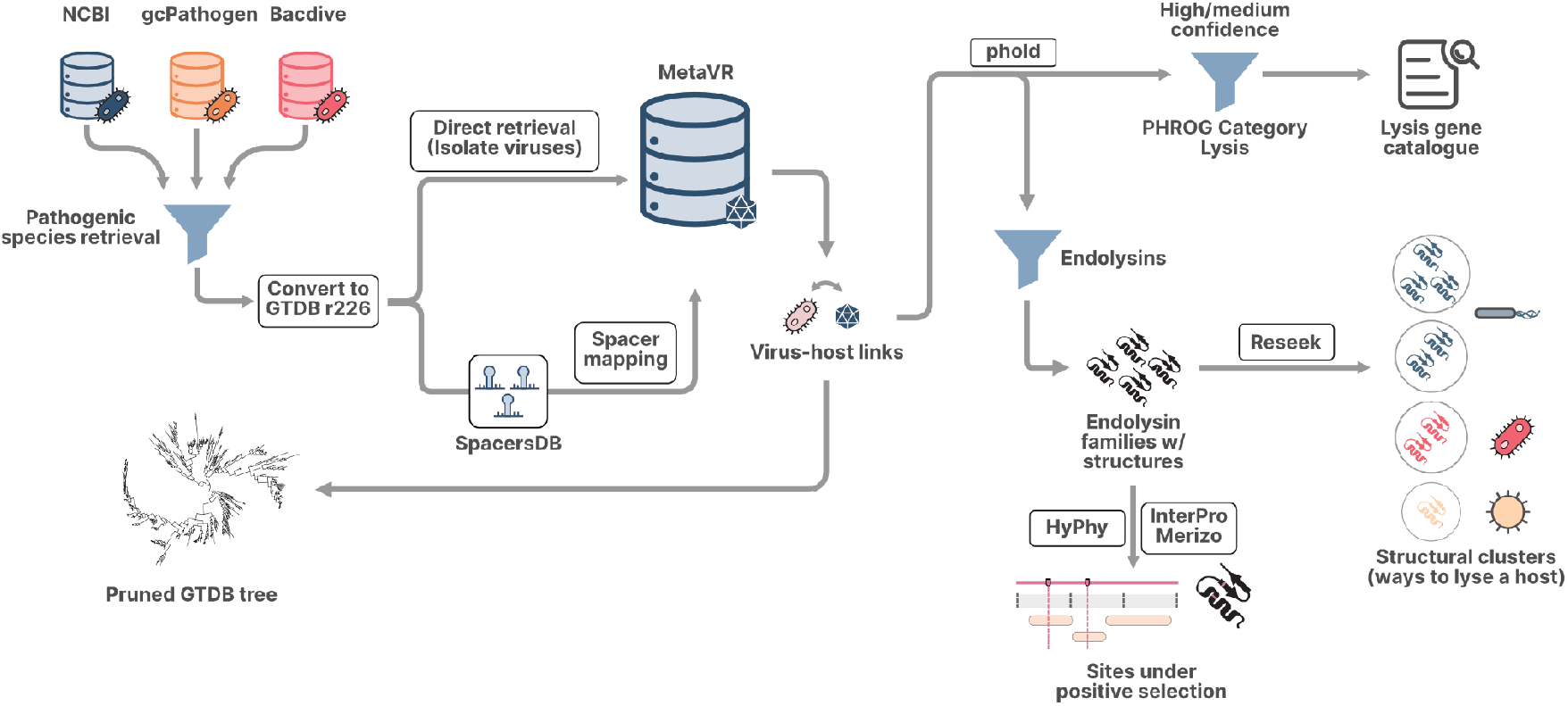

## Introduction

Antibiotics have transformed the modern world, with impacts on human health, agriculture, and animal husbandry. For instance, they have enabled large-scale operations in livestock production by mitigating the risk of disease outbreaks and increasing food security. For human health, the average lifespan has dramatically increased, in part due to better healthcare opportunities by providing effective treatments for fatal bacterial infections. However, despite these benefits, the misuse of antibiotics has contributed to the emergence and spread of resistant microorganisms, which now pose a major threat to both human and animal health. This has led into a global effort to search for alternative therapeutic strategies (1), such as the discovery of novel antibiotic molecules, the use of antimicrobial peptides that act by disrupting the microbial cell membrane (2, 3), the development of monoclonal antibodies for specific epitope binding (4), and exploration of secondary metabolites with antimicrobial properties from biosynthetic gene clusters (5).

These approaches, while promising and necessary, also have challenges associated that preclude their widespread availability as treatment. Antimicrobial peptides, for instance, often lack stability and can exhibit toxicity at therapeutic concentrations (6, 7), complicating their clinical application. Monoclonal antibodies on the other hand are constrained by complex manufacturing, risks of patient immunogenicity, stability issues, and prohibitive production costs (8, 9). Finally, the discovery pipeline for both novel small-molecule antibiotics and secondary metabolites is constrained by difficulties on functionally profiling newly identified compounds (5, 10). Collectively, these challenges underscore the need for various approaches to disease control.

An alternative strategy is bacteriophage (phage) therapy: the use of lytic viruses to specifically target and eliminate bacteria, particularly strains resistant to conventional antibiotics. Phage therapy offers several distinct advantages. Foremost among them is host specificity, with substantially less disruption of the broader microbiome than broad-spectrum antibiotics. Phages are also self-amplifying, replicating as long as the bacterial host is present. The clinical potential of this approach has been demonstrated in several case studies (11–13), both as a standalone therapy and in conjunction with antibiotics (14). Phages have also proven effective at disrupting bacterial biofilms (15), and strategies to create efficient phage cocktails that target multiple strains to broaden their therapeutic spectrum have been discussed previously (16).

Despite its clinical promise, the identification of optimal viral candidates for specific pathogenic strains remains a bottleneck for phage therapy, a non-trivial challenge given the typical narrow host range of most phages. Traditionally, discovery relies on enrichment-based isolation, where environmental samples are cultured alongside target pathogens to identify lytic activity (17). This culture-dependent workflow is both laborious and time-consuming, and frequently skews discovery toward fast-growing, highly prevalent pathogens.

Conversely, advancements in high-throughput sequencing and viromics have resulted in expansive viral databases (18–23) that create a new paradigm in phage discovery (24). These resources facilitate an accelerated hypothesis-testing loop by enabling the comparison of many more viral sequences and more efficient prioritization of candidates for functional validation across a broader host-range spectrum. However, while computational host-prediction tools such as iPHoP (25), CHERRY (26), and RaFAH (27) automate the linkage between phages and their putative hosts, the field is still in the early stages of establishing these connections with high confidence, often limiting predictions to the genus or family level. Consequently, the predictive power of genomic analysis remains constrained by a lack of species-level resolution and systematic, high-confidence linkages between these uncharacterized phages and their specific bacterial targets. As such, genomics-driven strategies for the identification and prioritization of therapeutic phages represent an important, yet underexplored, area of research.

In this work, we present a broad analysis of the MetaVR database (23) to identify phages that infect human bacterial pathogens. By mining this large-scale repository of viral genomes and augmenting its host predictions with CRISPR-spacer evidence from the JGI CRISPR-spacer DB (28) and SpacersDB (29), we established putative phage-host connections at the species level. Our analysis linked 196,472 high-quality and complete virus genomes representing 42,360 viral operational taxonomic units (vOTUs) to 618 species of human bacterial pathogens and opportunistic bacteria, substantially expanding the known diversity of viruses targeting clinically relevant microbes. Furthermore, we investigated the lytic genes encoded within these viral genomes and the predicted structures of their associated protein families. Focusing on endolysins, we identified potential enzybiotic candidates and explored the different evolutionary strategies employed by these phages for microbial cell degradation. Taken together, our results characterize phage diversity through functional, structural and evolutionary analyses, with potential applications to human health.

## Materials and Methods

### Compilation and classification of human bacterial pathogens

The NCBI pathogens (available at https://www.ncbi.nlm.nih.gov/pathogens/isolates/) resource was mined to identify bacterial species associated with human clinical isolates by filtering for isolates from *Homo sapiens*. NCBI Taxonomic assignments were converted to GTDB (30) r226 taxonomy by mapping against the Biosample ID on the GTDB metadata table. Additionally, the gcPathogens(31) database and Bacdive (32) were mined for additional human pathogens. Pathogen names from these two databases were converted to GTDB r226 using the taxopy v.0.14.0 package (https://github.com/apcamargo/taxopy). In total, we obtained 904 putative human pathogen bacteria species for further processing.

Predicted hosts were programmatically tagged according to the WHO Bacterial Priority Pathogens List (2024), available at https://www.who.int/publications/i/item/9789240093461. When a species matched multiple categories due to rank generalization, we assigned the pathogen to the category where its classification was most specific (for instance, *Escherichia coli* could fall under Critical as an Enterobacterales member or High as an *Escherichia member*, so it was assigned to the High group due to a more specific classification). Due to the lack of strain-level resolution when normalizing taxonomy via GTDB, antibiotic resistance data was not factored into the classification, instead, we focused exclusively o the broader taxonomic groupings to ensure a uniform comparison across the retrieved pathogenic species. WHO categories were thus defined as follows:

- Critical: All members of the order Enterobacterales (excluding those specified in the High Priority group as explained above), *Acinetobacter baumannii*, and *Mycobacterium tuberculosis*.
- High: Members of the genera *Escherichia* and *Salmonella*, alongside *Enterococcus faecium, Pseudomonas aeruginosa, Neisseria gonorrhoeae*, and *Staphylococcus aureus*.
- Medium: *Streptococcus pyogenes, Streptococcus agalactiae*, and *Haemophilus influenzae*.

One limitation of this programmatic classification is potential over-representation due to the low specificity of certain categories. For instance, the WHO Critical list includes the order Enterobacterales, a broad group that encompasses both pathogens and commensal/opportunistic species. To mitigate the possibility of flagging non-priority species, we focused most of our discussion on the most established clinical isolates.

### Retrieving viral genomes with host evidence

MetaVR includes host predictions generated from two methods: direct host associations (a virus detected from isolated organisms, or with verified hosts through VirusHostDB (33)) and computational host prediction generated using iPHoP (25), which aggregates and ranks the results of different computational host prediction tools. We mined the MetaVR database for viruses with direct evidence of infecting these 904 pathogens. Only genomes of High-quality or putatively complete viruses were considered (i.e. having direct or inverted terminal repeats topology, an estimated completeness greater than 90 or Quality as either “Complete” or “High-quality” by CheckV (34)). Viruses were classified as Isolate or Meta according to their source dataset: “Meta” if the dataset came from “Metagenome-assembled genome”, “Metagenome” or “Metatranscriptome”, and “Isolate” for all other cases.

Next, we sought to retrieve computational prediction results, however, iPHoP’s most specific assignment only goes to the genus level, which was not suitable for the analysis we were interested in this work. Thus, we computed host association for all retrieved viruses, including those with direct association, by searching for evidence of host interaction through CRISPR spacers on the pathogen genome.

We used two databases for this analysis: the CRISPR spacers database (28) and the SpacersDB (29). For the latter, spacers were filtered to be of high quality, and originating from genomes of high or medium confidence. Spacers from the human pathogen species of interest were kept and searched against all the complete and high quality viral sequences in MetaVR using spacerextractor (https://code.jgi.doe.gov/SRoux/spacerextractor). Alignments were retained when they contained at most one mismatch and covered at least 25 nucleotides. Identical spacer sequences were collapsed after treating each sequence and its reverse complement as equivalent. Associations supported by at least 10 distinct spacer sequences were retained as putative host links. In total, considering direct association and spacer assignment, we retrieved 196,472 virus genomes linked to 618 pathogens. The full table of host associations can be found in Supplementary table 1.

### GTDB tree retrieval and visualization

The GTDB r226 tree was pruned with ete4 v.4.3.0 (https://github.com/etetoolkit/ete4) to keep only the final 618 pathogen species, using *Peribacter riflensis* as an outgroup for rooting purposes. The same tree was also pruned to keep only species with at least 50 phages assigned for improved visualization in Figure 1. Trees were visualized with iTOL (35) and decorated with additional metadata such as WHO priority, gram-staining, number of vOTUs connected to the pathogen species, viral origin and host prediction method.

**Figure 1.**
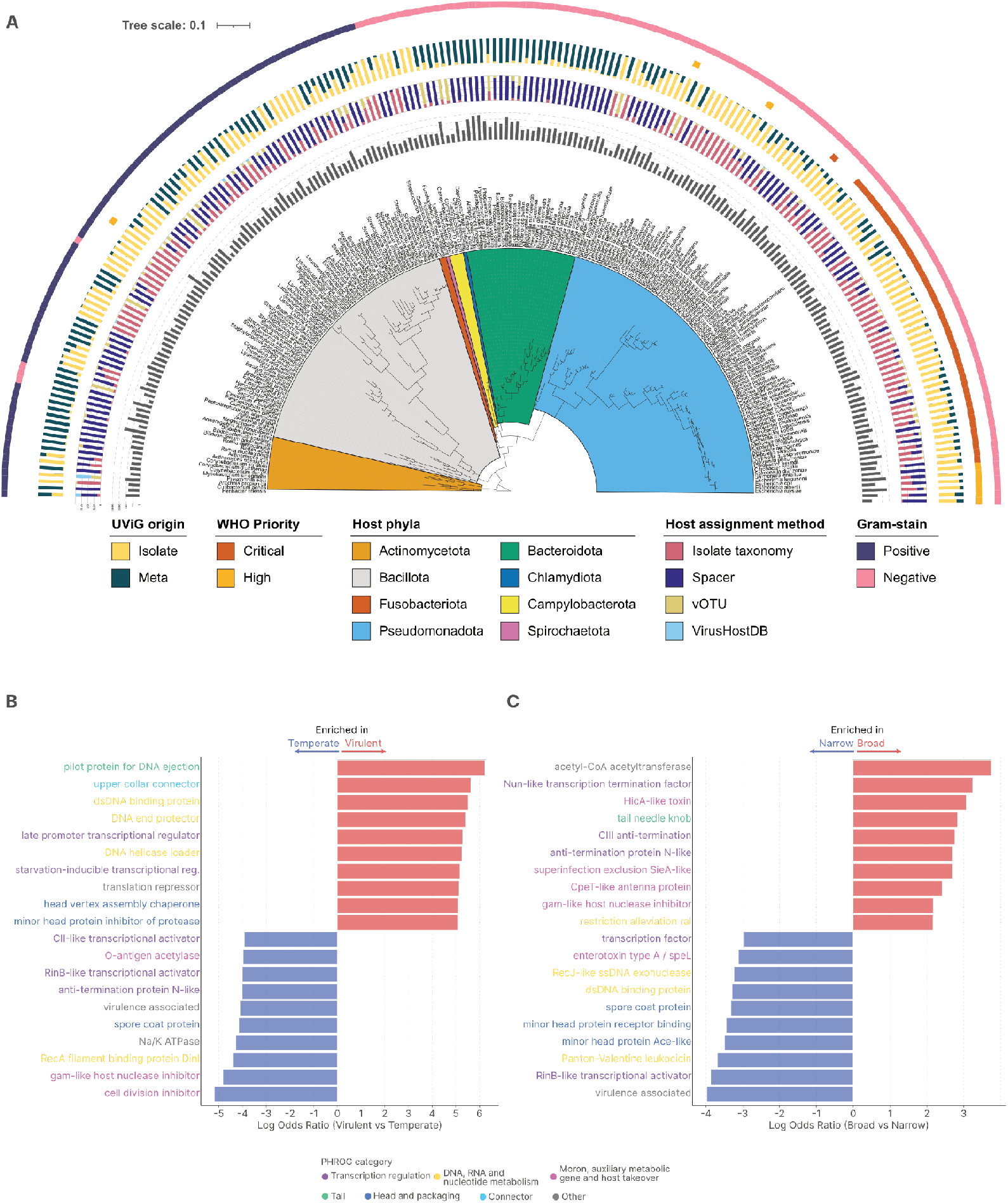
**(A)** GTDB r226 bacteria tree pruned at the species level to show only pathogenic bacteria with detected phages from MetaVR, with Peribacter riflensis as outgroup. Only species with at least 50 phages are shown for clarity (see Supplementary Figure 1 for full tree). The tree was decorated using iTOL with the number of vOTUs assigned to each species (innermost ring), the proportion of host assignment methods (second ring), the proportion of UViG origin (third ring), if the species is considered a WHO Critical or High priority pathogen (fourth ring) and the gram-stain of the species (outermost ring). Clades are colored according to their phyla. Top 10 most enriched Phold functions associated with phages with temperate or virulent lifestyles **(B)**, and broad or narrow host ranges **(C)**Phold functions are further colored according to their predicted PHROG category. Enrichment was calculated as the log of the odds-ratio from Fisher’s exact test (Benjamini-Hochberg adjusted p-values).

### Retrieving lysis genes

To identify the lytic potential within our viral dataset, we performed functional annotation using phold v.1.1.0 (36), filtering for high and medium-confidence predictions belonging to the ‘lysis’ PHROG category. These candidates were then mapped to their protein families, retrieved from MetaVR, to assess conservation and availability of a predicted structure representing them. We note that this approach was carried out even for prophages, allowing us to identify promising lysis-related across the whole dataset.

Lysis protein families with structural annotation were retrieved and their structures were subjected to an all-versus-all comparison using Reseek (37) (parameters: -dbsize 1049 -columns std+aq -sensitive). Structure pairs were then clustered according to the framework of Camargo et al. (2025) (19), by first retaining only structure pairs with bidirectional coverage greater or equal to 90% and alignment quality greater or equal to 0.85. These filtered pairs were then clustered using pyleiden v.0.1.2 (parameters: -r 0.44).

Structural clusters from the previous search where filtered to only have members from Meta sources, and the underlying structures were searched against the BFVD (38), VAD (39), PDB (40) and SCOP40 (41) structural databases, again using Reseek (parameters: -dbsize 1000000 -sensitive -columns std+qcovpct+tcovpct). Results were then filtered for p-value smaller than 0.05 and percent identity higher than 50%. Structures that did not pass these filters and with an average pLDDT over 70 were considered “novel” lysis-related candidates.

### Enrichment analysis and group comparison

We segmented our dataset into distinct groupings for analysis. The first grouping separated phages that were detected in “Isolate” sources (defined in MetaVR as coming from RefSeq, isolated bacteria or from single-cell sequences) and “Meta” sources (phages from metagenomes and metatranscriptomes). The second grouping referred to phages from “Broad” host range (vOTUs encompassing phages connected to 2 or more host genera) and “Narrow” host range (vOTUs encompassing phages connected to only species, or multiple species from a single genera). Finally, we also did a third grouping, by segmenting phages into “Virulent” and “Temperate” vOTUs by first running PhaTYP (42) via PhaBOX v2.1.13 against all phage genomes. Then, when collapsing into the vOTUs, we considered the results to be “Virulent” if all phages within the vOTU had a virulent assignment and there were no prophages in the vOTU. For all other cases the vOTU was considered as “Temperate”.

Enrichment of functions between Broad and Narrow host-range vOTUs, and Temperate and Virulent vOTUs was done through Fisher’s exact test with BH correction of p-values, as implemented in scipy v1.17.1. The number of vOTUs carrying a phold annotation against the number of vOTUs not carrying the same annotation was used for the contingency table. All results from this analysis can be seen in Supplementary table 2.

### Analysis of endolysins and comparison with the PhaLP 2.0 database

In addition to the broad exploratory analysis of lysis-related genes, we focused separately on characterizing the predicted endolysins. To investigate the structural diversity of our dataset, which could indicate different strategies phages employ to degrade the host cell wall, we again conducted an all-versus-all structural comparison of all endolysin protein families with available structural predictions using the same Reseek and pyleiden parameters as for all lysis-related protein families.

To evaluate how our endolysin sequences compare to existing lysin diversity, we performed a comparative analysis against the PhaLP 2.0 database (43), a large database of predicted lysins obtained from UniProt and EnVhog. Following their workflow, sequences from both sources were clustered via MMSeqs2 (44) v. 17-b804f (parameters: –min-seq-id 0.3 -c 0.7).

Enrichment analysis of species connected to MetaVR exclusive against shared with PhaLP 2.0 endolysin clusters was also done with Fisher’s exact test with BH correction of p-values.

### Selection analysis and domain annotation

To identify the evolutionary drivers of endolysin evolution, we performed a comprehensive selection analysis on the protein families, searching for evidence of positive selection on sites that could be involved in the lysis process. For each protein family with a structural prediction, coding sequences (CDS) were extracted using pyrodigal-gv (45). Multiple sequence alignments were generated with FAMSA2 (46), which subsequently were used for phylogenetic inference via FastTree2 (47). Codon-aware alignments were then produced by back-translating the CDS from the protein alignments using PAL2NAL (48). Finally, we assessed pervasive and episodic diversifying selection using HyPhy FUBAR (49) and MEME (50), respectively. This entire workflow was implemented in a Nextflow pipeline to streamline processing.

Domain identification and segmentation were performed on the representative proteins for each family under selection using InterProScan6 (51) and Merizo (52). We further discuss (Figure 4) families with high-quality structural prediction where selection sites fell within domains found by both sequence and structural methods. Selection analysis results for all evaluated families are provided in Supplementary table 4.

## Results and Discussion

### Genome-scale mining links thousands of phages to human bacterial pathogens

We were able to link 196,472 high-quality and complete phages (95,034 from isolate sources, 101,438 from meta sources) representing 42,360 vOTUs to 618 human-pathogenic or opportunistic bacteria (Supplementary Figure 1, Supplementary table 1). From these vOTUs, we found 10,241 vOTUs comprised of lytic phages, 6,817 of which did not have any phage sequences classified as lysogenic. A large amount (99,315) of detected phage sequences were prophages, with 24,260 vOTUs (6,564 non-singletons) being exclusively comprised of prophages. Although temperate phages are generally less suitable for direct therapeutic use, their genomes may nevertheless encode lytic proteins with potential therapeutic applications.

Figure 1 highlights species with at least 50 phages connected to them. Most phages associated with a World Health Organization (WHO) Critical priority were from isolate sources, which could indicate that their hosts have been sequenced more often, while the majority of phages from Bacteroidota, Campylobacterota and Actinomycetota were found in Meta sources. Most of the organisms found exclusively or predominantly in Meta sources can be characterized as opportunistic pathogens. Some examples include members of the genera *Bacteroides* and *Prevotella* in the gut and Rothia in the oral cavity, which are typically stable components of the human microbiome but can cause disease if they move from their primary body sites. Notably, our mining strategy recovered predominantly Meta-derived phages for opportunistic pathogens previously represented by few isolate members. For example, *Leminorella richardii* was linked to 52 UViGs (29 vOTUs), with 48 UViGs originating from Meta sources. *Leminorella* species have been reported as multidrug-resistant nosocomial pathogens associated with UTIs, surgical-site infections, bacteremia and other infections (53).

As expected, most vOTUs were linked to a single species, with a sharp decline in the number of vOTUs linked to 2 or more species (Supplementary Figure 2). We further explored if there would be any functional evidence dictating host specificity, seeking to understand why certain phages are able to infect multiple genera (broad host range), while others are restricted to a single species or a few species on the same genera (narrow host range), and possible relations with viral lifestyles. We also explored how these functional determinants correlate with the phages’ distinct evolutionary lifestyles.

Comparison between virulent and temperate phages revealed functional enrichments that reflect their different strategies (Figure 1 b). Virulent phages were strongly enriched in structural and replication-associated functions, as illustrated by half of the top 10 terms being related to head and packaging or nucleotide metabolism, consistent with a strategy of rapid genome replication, particle assembly and host lysis. An interesting observation is a lysis inhibition product (ranked 11th, log-odds=5, p_adj<0.001), as it has been reported that phages such as T4 delay lysis of the host cell when detecting secondary infections as a way to optimize progeny production (54, 55). Temperate phages however showed enrichment for transcriptional regulators and host-interaction factors, such as the CII- and RinB-like regulators, anti-termination proteins and anti-RecBCD systems. These functions indicate a strategy of controlling gene expression and modulating host defenses to allow long term stability within the host genome.

The distinction between broad and narrow host-range phages was less defined (Figure 1 c), but still revealed interesting trends. Broad host-range phages were enriched in functions associated with overcoming host defenses, including anti-RecBCD systems and the Restriction alleviation protein (Ral). Narrow host-range phages, on the other hand, were enriched in transcriptional regulators and factors associated with specialized host-virus interactions. These include receptor-binding components and host modulation factors, such as the Enterotoxin type A product, which has been reported to be encoded by a Staphylococcal phage and important for the toxicity phenotype of the host (56, 57). The presence of phage-encoded enterotoxins indicates a strategy where the phage enhances host fitness by tightly coupling it to its own genome, ensuring long-term persistence. These results suggest that narrow host-range phages prioritize niche-specific adaptation, utilizing tight regulatory coupling and fine-tuned metabolic integration with their specific hosts to maintain stability.

Interestingly, many functions enriched in temperate phages overlap with those considered as broad host-range. Some examples include anti-defense systems, anti-termination proteins and transcriptional regulators. These shared modules suggest a generalized evolutionary strategy to improve compatibility across different bacterial groups. By deploying these tactics, the phage can bypass host immunity and hijack regulatory networks, creating stability for its long-term maintenance via the lysogenic cycle.

Taken together, these results demonstrate that the phage functional architecture is tied to its ecological and evolutionary lifestyle. While virulent phages are enriched in functions supporting rapid replication and efficient lysis, the success of both temperate and broad host-range phages frequently depends on subverting bacterial immunity or conferring certain fitness advantages to their hosts. This relation between viral persistence and bacterial adaptation can be a challenge for whole-phage therapeutics, as many broad host-range phages carry cargo genes that can benefit the host, hindering their direct clinical use without prior genomic modification. However, these findings also underscore the value of targeted genomic mining; although temperate phages may be unsuitable for direct therapeutic application, their genomes represent a potentially valuable reservoir of lysis-associated proteins that can be exploited independently of the intact phage. We therefore next characterized the lysis-associated gene repertoires of the detected phages.

### Meta-derived phages reveal structurally convergent yet sequence-diverse lysis machinery

To determine if viruses from Meta sources would reveal novel lysis strategies distinct from those from cultivated isolates, we compared the distribution of lysis-related protein families across both datasets (Figure 2 a and b). While sequence-level analysis revealed a high proportion of exclusive families for both groups, this diversity largely converged at the level of predicted structures. The overlap in structural architectures between the Isolate and Meta datasets is suggesting that substantial sequence divergence in lysis-associated proteins frequently occurs within a more limited repertoire of structural architectures. Nevertheless, by clustering these 1,969 lysis-related families with structural prediction using Reseek (37) and pyleiden (see Methods), we identified 236 structural clusters composed exclusively of Meta sources. These clusters comprise 13 non-singleton groups (4 with at least 3 members) and a total of 256 individual structures. We then took these 256 structures and searched against the BFVD (38), VAD (39), PDB (40) and SCOP40 (41) structural databases to identify folds with low structural similarity. By defining “known” structural homologs as those with a p-value lower than 0.05 and identity of at least 50%, we found that 76 structures failed to meet these criteria despite having high-confidence (pLDDT > 70) structural predictions (Figure 2 c).

**Figure 2.**
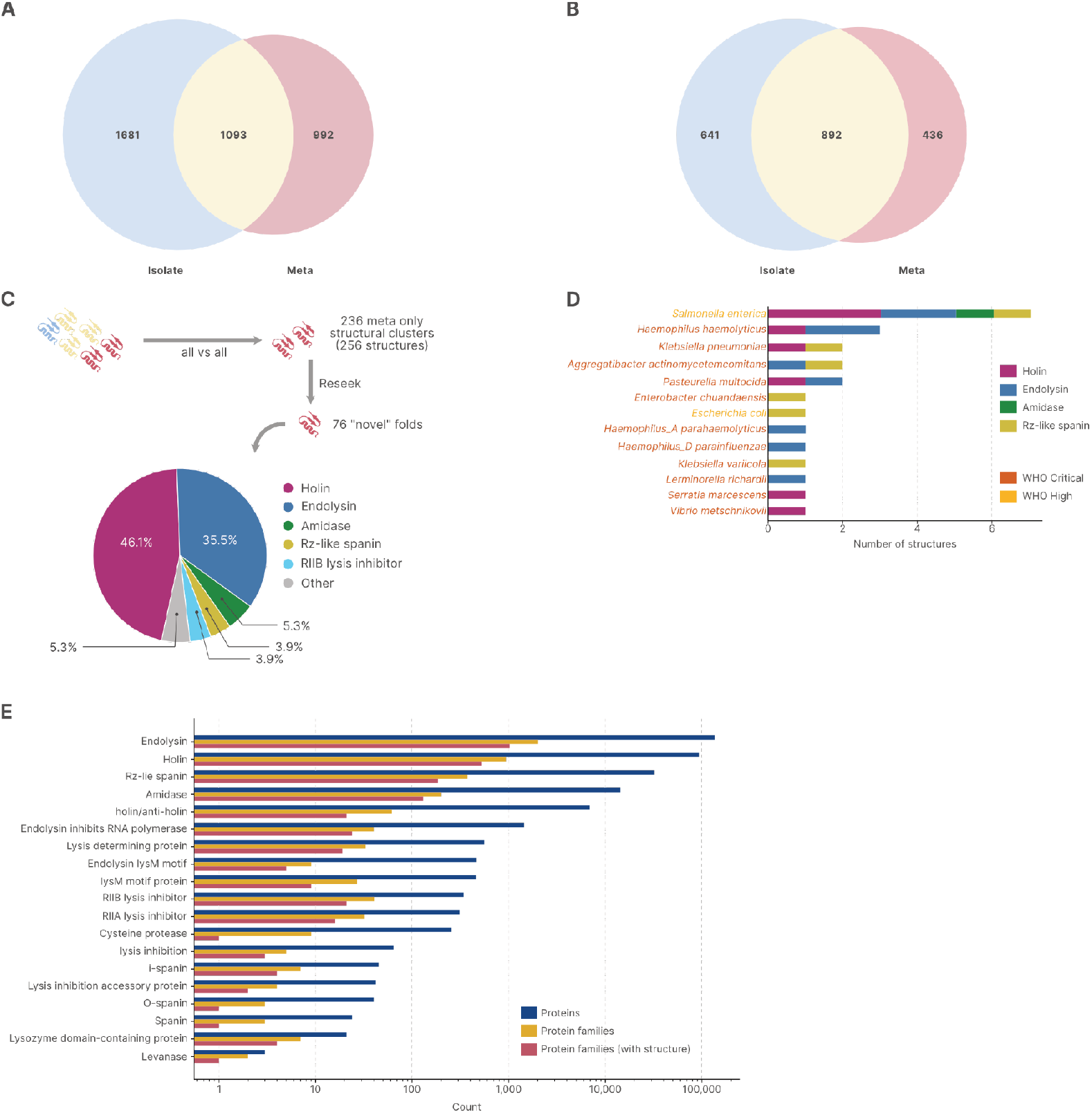
Number of lysis protein families **(A)** or lysis protein families with structural prediction **(B)** with members exclusively from phages from meta sources, isolate sources, or shared between both. **(C)** Schematic of search for novel lysis related structures and distribution of phold annotations of novel structures. **(D)** WHO high and critical priority pathogens targeted by the “novel” structures. **(E)** Number of proteins, protein families and protein families with structures for each lysis-related predicted annotation

We next determined if any of these 76 structures were associated with WHO High or Critical priority pathogen species and identified associations with 13 pathogen species (Figure 2 d). These results suggest a reservoir of novelty within the virosphere, representing putative novel lysis enzymes that could serve as candidates for further investigation as enzybiotics against clinically relevant pathogens. This structural novelty is particularly relevant given that many bacteria have evolved sophisticated mechanisms to evade lytic enzymes. For instance, it has been reported that some bacteria modify peptidoglycan cross-links to hinder endolysin interaction (58), while others, such as *Listeria monocytogenes* and *Staphylococcus aureus*, decorate the peptidoglycan backbone with molecules to escape detection (59, 60). Indeed, we found two endolysin structures, one amidase and one spanin, from this “novel” structures group putatively targeting *L. monocytogenes*. The discovery of these structures targeting a pathogen known for its complex cell-wall modifications further strengthens the hypothesis that Meta-derived viromes harbor untapped lysis-associated diversity that could be capable of overcoming previously challenging bacterial defense mechanisms.

The largest group of lysis-related genes were endolysins (Figure 2 e), followed by holins and spanins. Holins act with endolysins by forming pores in the inner cell membrane to allow endolysins to access the peptidoglycan layer. Spanins, on the other hand, are responsible for disrupting the outer membrane of gram-negative bacteria, allowing the release of viral particles (61). The fact that most lysis-related protein families with structural prediction were endolysins could indicate that they are more conserved and thus more likely to be structurally predicted by grouping in sufficiently large families, while holins and spanins might be more diverse and thus in smaller protein families.

### Structural diversity of endolysins reveals shared repertoires across pathogen species

As we identified several endolysins in the phages linked to human pathogens, we further analyzed their structural predictions to understand evolutionary strategies that might be associated with microbial cell degradation. Specifically we were interested to know how many unique structures they would represent, as different structures could mean different strategies to lyse a given host. We thus retrieved the 1,126 structural predictions for each endolysin protein family connected to a pathogen and clustered them using Reseek (37) and pyleiden (see Methods), resulting in 592 structural clusters. From these, 397 were singletons, and the largest cluster had 16 members, indicating how diverse these structures can be.

We represented each pathogen by the set of endolysin structural clusters associated with its linked phages and calculated pairwise Jaccard similarity between these repertoires. Figure 3 reveals similarity among Enterobacterales, which shared a median of 15 structural clusters per species pair and has substantially greater repertoire similarity than non-Enterobacterales pairs. Strong within-genus similarities were also evident among *Klebsiella* and *Acinetobacter* species, including the ESKAPE (62) pathogens *K. pneumoniae* and *A. baumannii*. This pattern is consistent with related bacterial hosts being targeted by phages encoding structurally related endolysins. These shared structural families provide candidates for experimental testing of a cross-species lytic activity cocktail.

**Figure 3.**
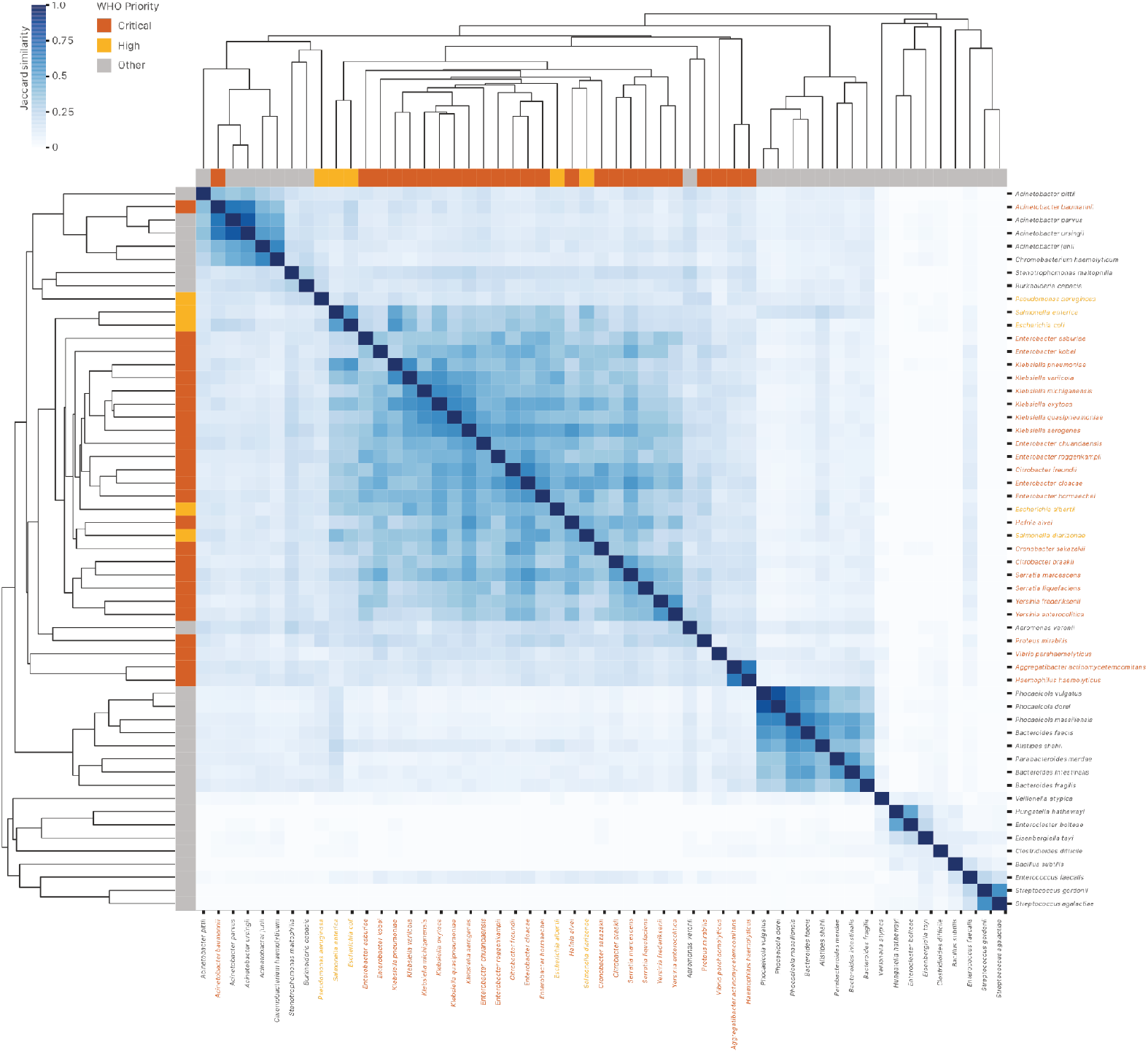
Hierarchically clustered heatmap of pairwise Jaccard similarity in phage endolysin structural-cluster repertoires across pathogen species. Jaccard similarity between two species was calculated from the overlap in their endolysin structural clusters. Species are clustered using average linkage on the pairwise distance matrix (1 – Jaccard). Side color bars indicate WHO priority category (Critical, High, or other). Only species connected to at least 20 distinct structural clusters are shown for clarity.

To contextualize our findings, we also compared the endolysin proteins identified in this study against the PhaLP 2.0 database (43), a comprehensive repository of endolysins predicted using the sublyme tool. By clustering our predicted sequences with the PhaLP 2.0 endolysins following their parameters (Methods). We observed that while the majority of our endolysins were assigned to shared groups with their sequences and, interestingly, the number of total clusters reduced slightly from 25,587 to 25,401, indicating sequences from this work bridged the gap between a few separate PhaLP clusters. Additionally 380 clusters (13,255 sequences) were unique to this study (Supplementary Figure 3 a), with three of them having more than 1,000 endolysins (Supplementary Figure 3 b) and 40 being comprised exclusively of Meta sequences (23 non-singletons). However, only 164 sequences of these 380 clusters were classified as endolysins by sublyme. This could reflect the algorithmic differences between how sublyme and phold annotate: while phold relies on structural homology, sublyme uses a machine-learning classifier trained on curated lysin sets, though a subset may represent genuinely novel endolysins absent from the training data.

Considering that the PhaLP 2.0 resource encompasses endolysins from phages across diverse bacterial lifestyles and is a much larger resource, these distinct clusters suggest a specialized repertoire of endolysins associated with human pathogens, potentially reflecting niche-specific adaptations in viral lysis machinery that could be leveraged for practical use-cases. To verify this claim, we compared species proportions between the exclusive and shared clusters using Fisher’s exact test (BH-corrected). While E. coli and S. enterica were proportionally represented in both groups, several clinically important species were significantly enriched in the exclusive clusters (Supplementary table 3), including *C. difficile* (log-odds=2.41, adj. p<0.001), *K. pneumoniae* (log-odds=1.13, adj. p<0.001), and *B. melitensis* (log-odds=5.16, adj. p<0.001).

### Positive selection has shaped the functional domains of several endolysin protein families

To determine whether evolutionary pressures might be driving enhanced catalytic efficiency, we investigated endolysin protein families for evidence of positive selection. Because advantageous amino acid substitutions could optimize cell-wall hydrolysis, we analyzed individual sites within these families for evidence of both pervasive (49) and episodic (50) diversifying selection. We identified sites under positive selection within 278 endolysin families. Of these, 212 exhibited episodic selection, 9 showed pervasive selection, and 57 displayed evidence of both. To functionally contextualize these evolutionary pressures, we annotated the representative sequence of each family (the sequence utilized for initial MetaVR structural prediction) at the amino acid level using InterPro (51). This analysis revealed that in 167 families, the sites under selection were within predicted functional domains. Furthermore, in five families, these positively selected sites mapped directly to highly conserved residues annotated by CDD or PIRSR. This direct selection on conserved sites suggests that evolutionary pressures are actively modifying core catalytic or binding interfaces, potentially driving changes in substrate specificity, structural stability, or overall enzymatic efficiency.

To connect these sequence-level findings with the protein 3D architecture, we analyzed these 167 families at the structure level using Merizo (52), which indicated 65 families with sites under positive selection within segmented domains. Corroborating sequence finding, 60 families showed positively selected sites within predicted domains from both InterPro and Merizo. From a clinical perspective, 118 of the 167 families were connected to pathogens from WHO High or Critical categories, including 32 exclusively connected to WHO Critical and 20 to WHO High, indicating a significant group of enzymes with medical relevance. Results for all families under selection can be viewed on Supplementary table 4.

Figure 4 highlights three representative endolysin families with high-quality structural models, illustrating sites under pervasive and episodic selection on domains annotated via InterPro and Merizo. Notably, all three families comprise endolysins derived from phages targeting WHO High or Critical priority pathogens, with family IMGVR_PC_000548983 (Figure 4 c) found to be exclusively associated with *Mycobacterium tuberculosis*. As shown in Figure 4 b and Figure 4 c the sites undergoing selection in these families are predominantly found within predicted enzymatically active domains. This localization suggests that evolutionary pressures have acted in these catalytic regions, possibly optimizing their hydrolytic efficiency to counter host adaptations. In Figure 4 d we observed positive selection acting upon an amidase domain. This aligns with existing literature on *Staphylococcus aureus* phage endolysins, which leverage amidase domains for better hydrolytic activity, and endolysin variants lacking these specific domains have been shown to exhibit reduced lytic efficacy (63). Indeed, this protein family occurred in 43 UViGs spanning 10 vOTUs linked to *S. aureus*, providing a well-supported example of positive selection acting on a clinically relevant endolysin family.

**Figure 4.**
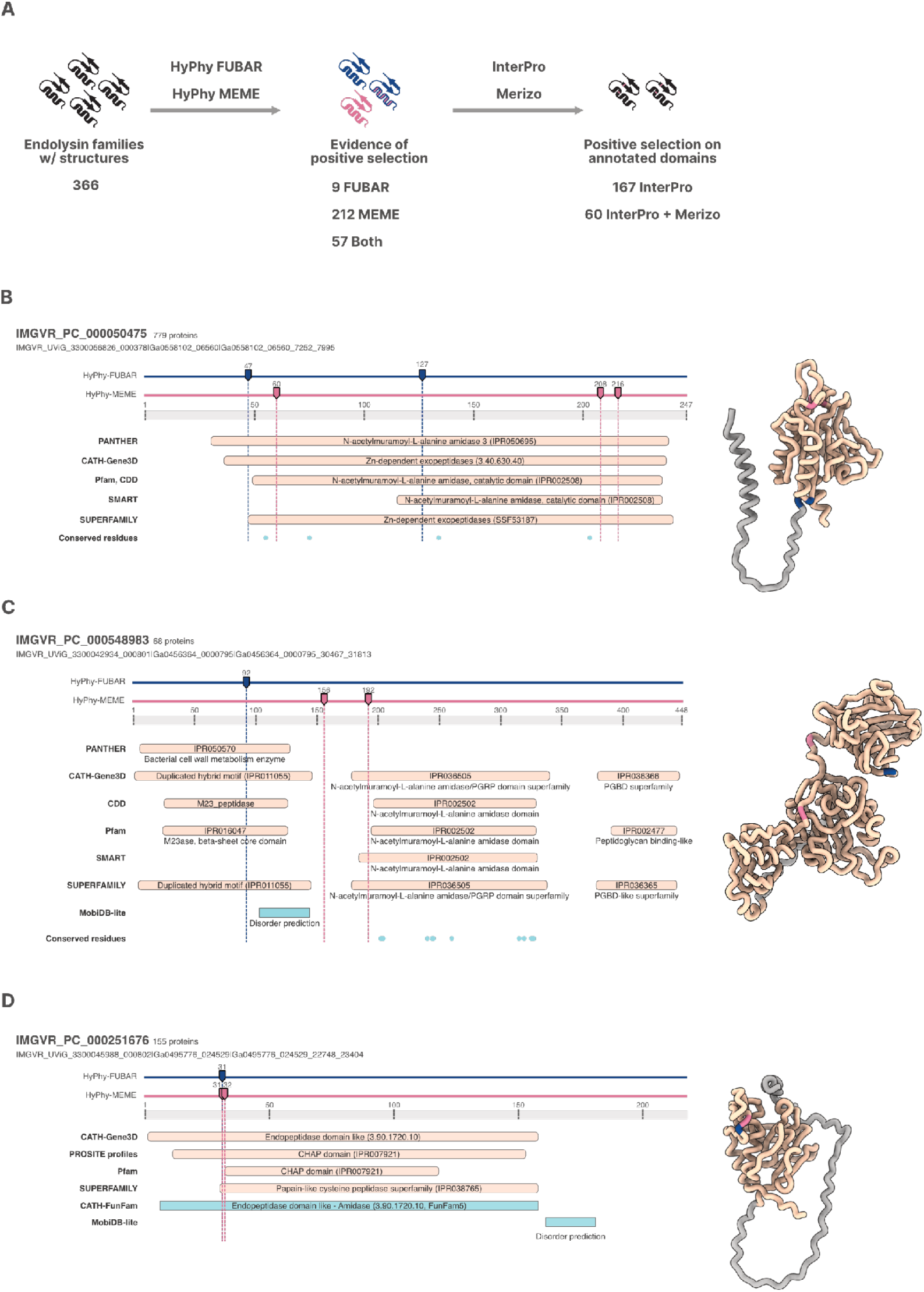
**(A)** Selection analysis workflow for endolysin protein families with structural prediction. **(B-D)** Schematic indicating site positions under positive selection for three endolysin families with evidence for both pervasive (FUBAR) and episodic (MEME) selection in functional domains predicted through sequence methods (InterPro) or structure segmentation (Merizo). Left-side figure shows the sites under selection with the InterPro predicted domains colored in yellow and regions/motif/transmembrane in blue, while right-side figures show the predicted 3D protein structure, with segmented domains colored in yellow. In both visualizations FUBAR sites are colored in dark-blue and MEME sites in pink.

Taken together, these results highlight how leveraging selection analyses, and the hypothesis that positive selection acting on protein domains, can guide the choice of candidates for further experimental investigation.

## Conclusions

In this work, we mined the MetaVR database to identify phages putatively connected to 618 species of human bacterial pathogens and opportunistic bacteria, linking 196,472 high-quality and complete viral genomes through direct host associations and CRISPR-spacer evidence. Functional enrichment analysis confirmed that virulent phages are enriched in functions associated with rapid replication, particle assembly and lysis, while temperate and broad host-range phages share strategies for subverting host defenses and modulating gene expression. By characterizing the lysis machinery encoded by these phages, we identified 76 apparent novel lysis-associated proteins a from structural stand-point, coming from Meta sources, with no close homologs in existing structural databases and including candidates targeting WHO priority pathogens. More focused analysis of endolysin protein families uncovered 592 structural clusters with extensive repertoire sharing among ESKAPE and other clinically relevant pathogens. Finally, selection analysis revealed 167 endolysin families with positively selected sites within functional domains, indicating that evolutionary pressures are actively reshaping catalytic and binding interfaces in these enzymes.

A few limitations should be considered when interpreting our results. First, the pathogen set may contain biases related to the host taxonomic normalization process. The GTDB classifications for instance, can obscure clinically relevant distinctions by collapsing pathogens with distinct clinical profiles due to genomic similarity, such as for *Shigella* species, which are considered *Escherichia coli* in GTDB. Additionally, the WHO priority mapping operates mostly at broad taxonomic ranks and for specific resistance phenotypes, which may inflate the number of species classified as Critical or High priority by including commensal or opportunistic organisms alongside established pathogens, as well as representative genomes in the CRISPR or isolate databases connected to a given phage not necessarily being an individual having the resistance phenotype. We attempted to mitigate this by focusing our discussion on the most established clinical isolates and by assigning species to the most specific applicable WHO category.

Second, CRISPR-spacer matches, while informative, do not constitute direct evidence of productive phage infection. A spacer may reflect an abortive infection, or result from the incorporation of environmental DNA fragments into the CRISPR array rather than a true phage encounter. We mitigated this risk by applying stringent filtering criteria, requiring a minimum alignment, almost no mismatches and a substantial amount of different spacer hits, which reduces but does not eliminate the possibility of spurious associations. Our results should therefore be interpreted as putative host-phage linkages that warrant experimental validation.

Third, while our computational analyses identify promising enzybiotic candidates, experimental validation remains essential. The 76 structurally “novel” lysis proteins require biochemical characterization to confirm their lytic activity, substrate specificity, and therapeutic potential. Similarly, the endolysin families under positive selection represent computational predictions of adaptive evolution; determining whether the identified substitutions confer functional advantages, such as enhanced lytic efficiency or altered host-range specificity, will require further assays. The high proportion of prophages in our dataset (99,315 sequences) also presents practical challenges for whole-phage therapeutic development, as converting temperate phages into lytic requires careful engineering to address superinfection immunity systems, toxin-antitoxin modules and the risk of horizontal gene transfer. However, these prophage-derived sequences are still relevant from an enzybiotic perspective by sidestepping the need for lifestyle conversion. Endolysins from prophages can be recombinantly expressed and engineered independently, making the large prophage fraction in our dataset an asset rather than a limitation.

Despite these limitations, our results establish a systematic, genomics-driven framework for prioritizing phage-derived therapeutics against clinically relevant bacteria and prioritizing phage-derived proteins for therapeutic investigation. By extending the search beyond cultivated phages, MetaVR reveals a large reservoir of lysis-associated sequence and structural diversity that remains largely unexplored. The identification of shared endolysin repertoires across ESKAPE pathogens provides a rational basis for designing broad-spectrum enzybiotic cocktails, while the catalog of novel lysis structures from Meta phages as well as the evidence of evolutionary pressures shaping enzymes expand the search space for candidates beyond what culture-based approaches have uncovered. Together, these results illustrate how large-scale viral genomics can transform the rapidly expanding sequence space of uncultivated viruses into a resource for discovering and prioritizing new therapeutic candidates against drug-resistant bacterial infections.

## Data and Code Availability

Phage sequences, protein families and protein structures are available in MetaVR (https://meta-virome.org). Code used for analysis is available at https://code.jgi.doe.gov/MBFiamenghi/uvigs_from_pathogens. All generated data is also available at Zenodo under 10.5281/zenodo.21970973.

## Acknowledgements

This work was conducted by the US DOE JGI (https://ror.org/04xm1d337), a DOE Office of Science User Facility, supported by the Office of Science of the US DOE operated under contract no. DE-AC02-05CH11231. A portion of this work was also supported by the US DOE, Office of Biological and Environmental Research (BER) as part of BER’s Genomic Sciences Program (GSP) under FWP 70880. M.B.F was supported by NIH 1U01DE034196-02 and NIH 1U01DE034196-03.

## Supplementary figures

**Supplementary Figure 1.**
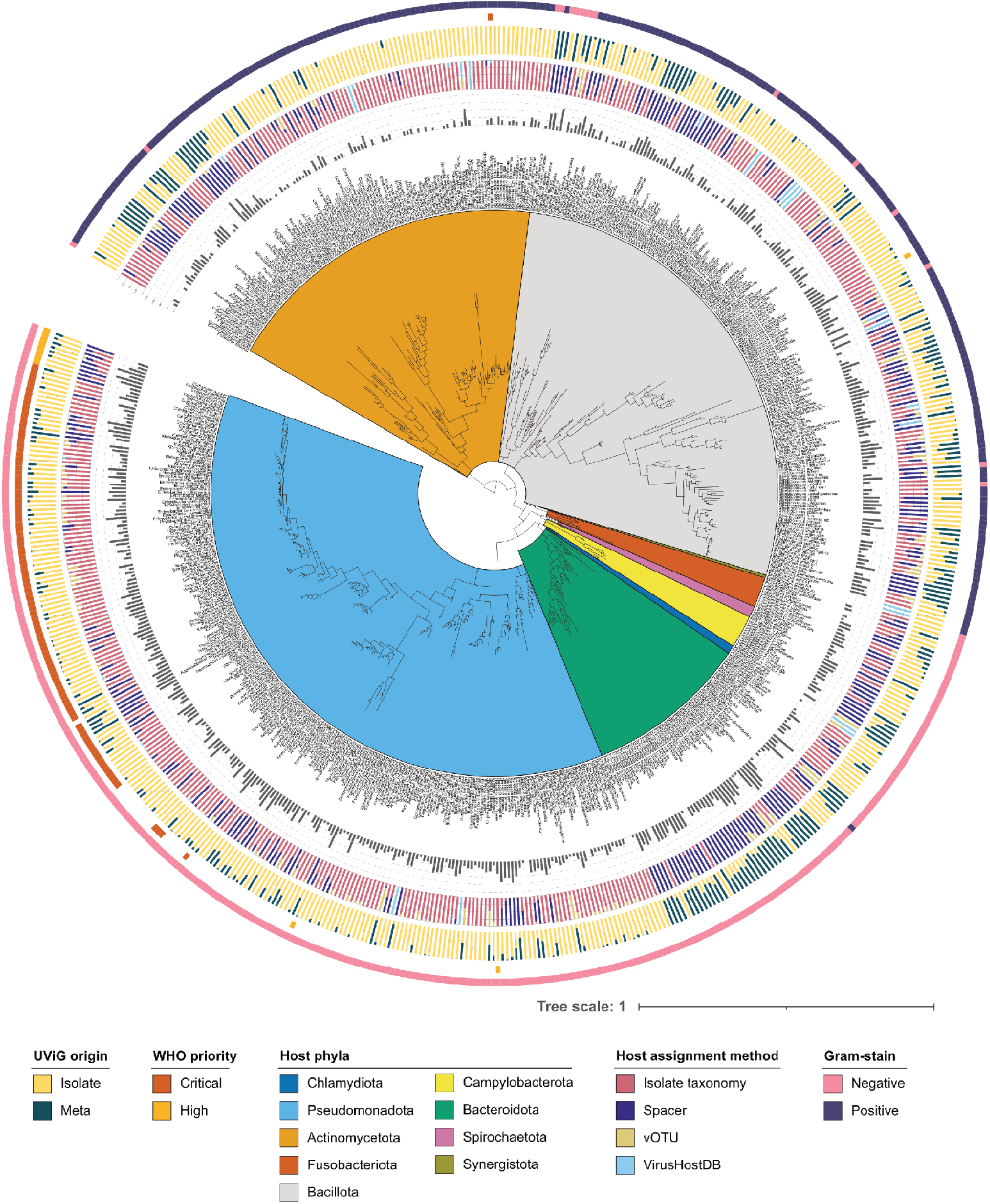
GTDB r226 bacteria tree pruned at the species level to show only pathogenic bacteria with detected phages from MetaVR, with Peribacter riflensis as outgroup. The tree was decorated using iTOL with the number of vOTUs assigned to each species (innermost ring), the proportion of host assignment methods (second ring), the proportion of UViG origin (third ring), if the species is considered a WHO Critical or High priority pathogen (fourth ring) and the gram-stain of the species (outermost ring)

**Supplementary Figure 2.**
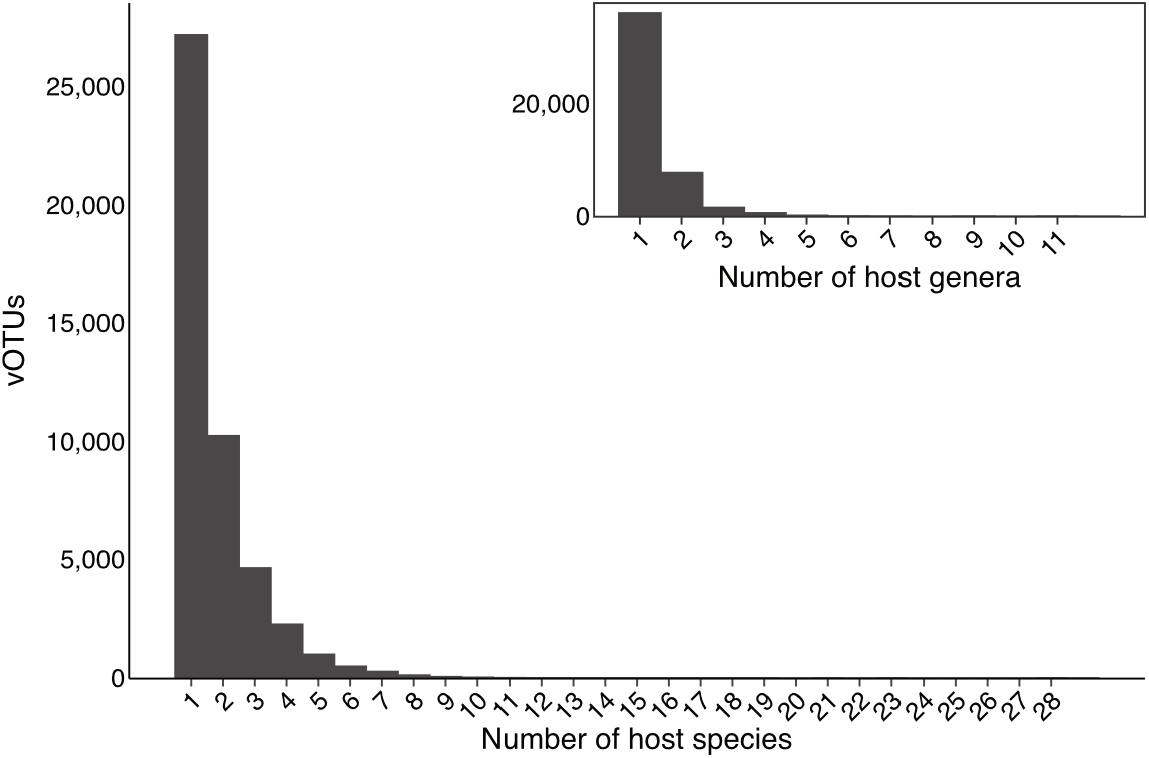
Number of different pathogenic species and genera (inset graph) hosts connected to each vOTU. Most vOTUs were assigned to a single species, although a non-negligible amount had broader host ranges

**Supplementary Figure 3.**
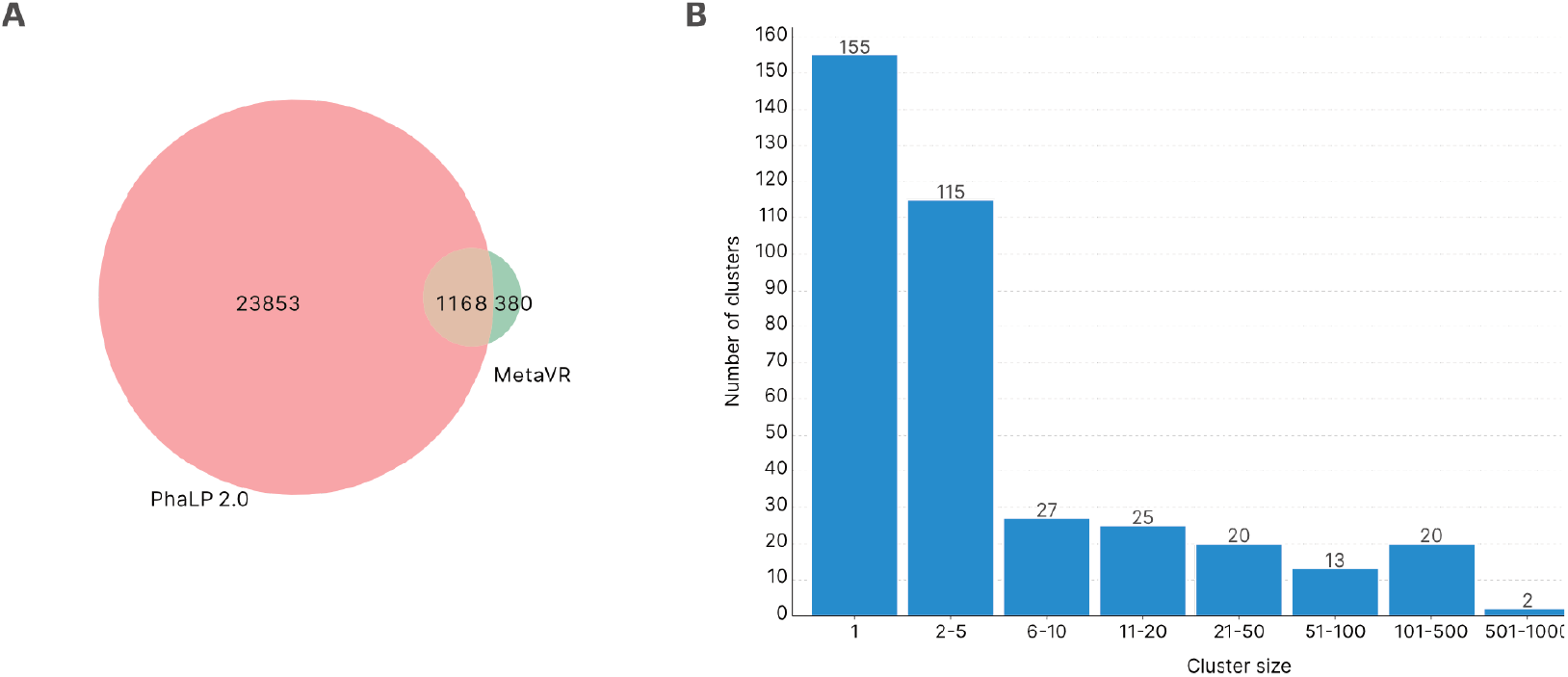
**(A)** Number of protein families generated when clustering endolysins from this work with the PhaLP 2.0 database. **(B)** Size distribution of the 380 generated clusters exclusive to this study

