## Supplementary figures and images for "Discovery of novel enzybiotic candidates targeting human bacterial pathogens through large-scale viral-host profiling"

### Supplementary Figure 1

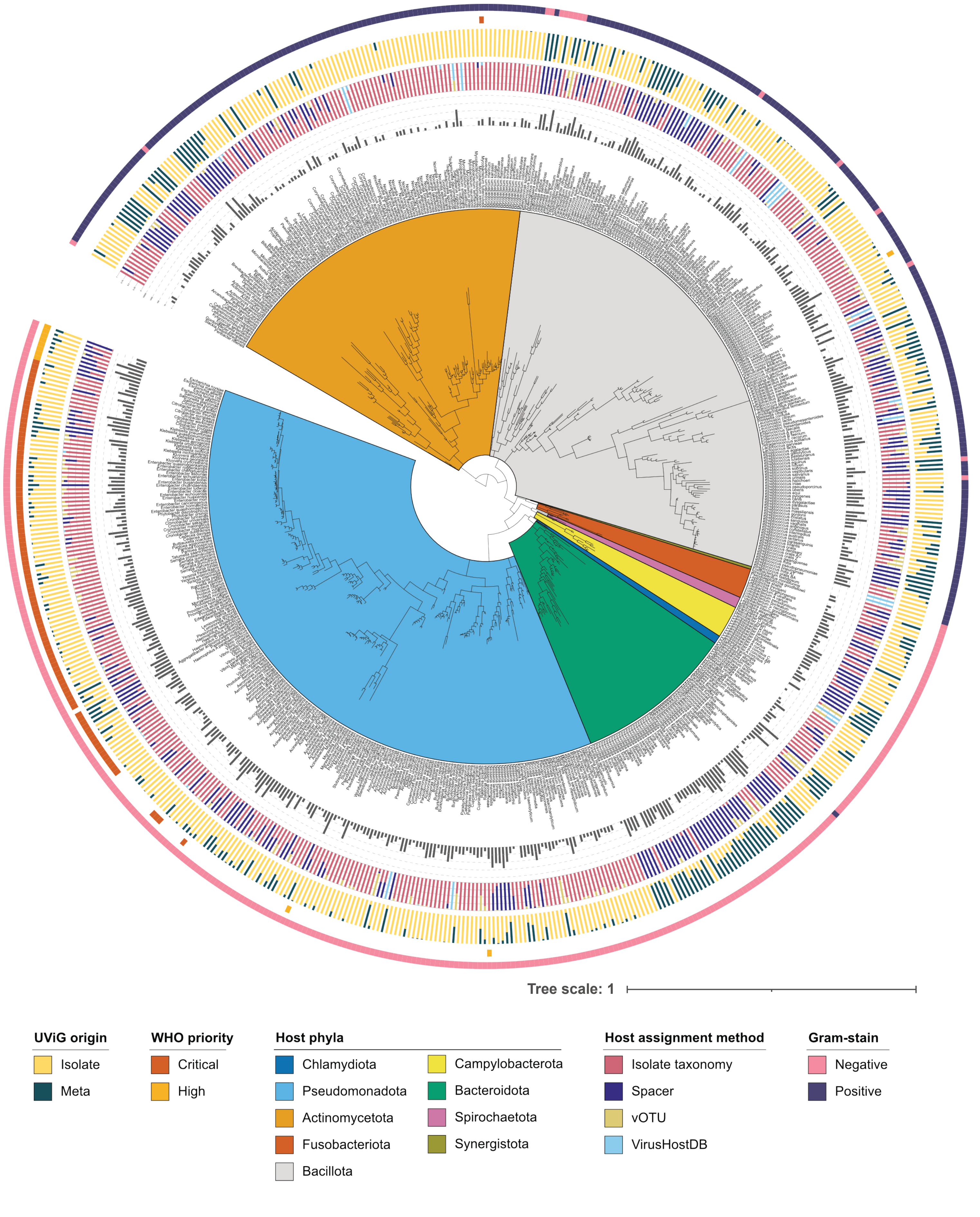

### Supplementary Figure 2

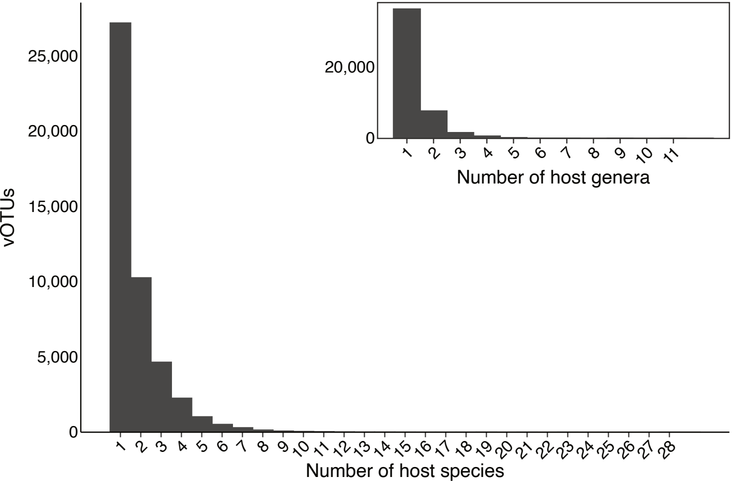

### Supplementary Figure 3

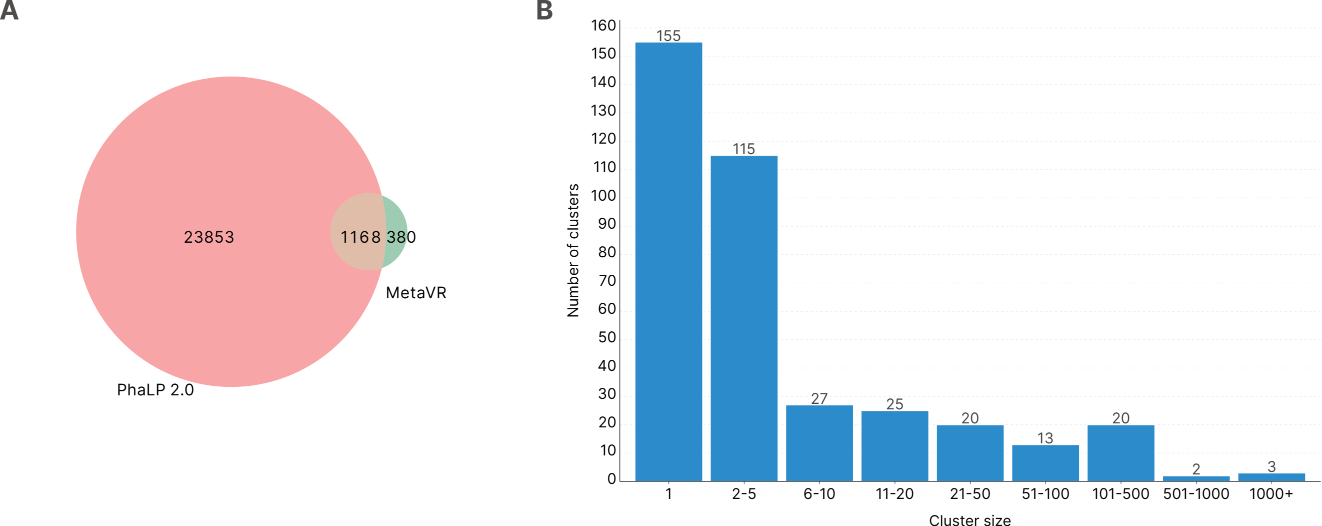
